# Flight behaviour diverges more between seasonal forms than between species in *Pieris* butterflies

**DOI:** 10.1101/2023.11.27.568806

**Authors:** Irena Kleckova, Daniel Linke, Francisko De Moraes Rezende, Luca Rauscher, Camille Le Roy, Pável Matos-Maraví

**Affiliations:** Institute of Entomology, Biology Centre CAS (Czech Academy of Sciences), České Budějovice, Czechia; Department of Zoology, Faculty of Science, University of South Bohemia, České Budějovice, Czechia; Experimental Zoology Group, Wageningen University, Wageningen, the Netherlands

**Keywords:** Flight kinematics, trajectory tracking, Lepidoptera, morphometrics, polymorphism

## Abstract

In flying animals, wing morphology is typically assumed to influence flight behaviours. Whether seasonal polymorphism behaviour might be associated with divergent fore wing shapes. *Pieris* individuals in the spring are small and have elongated fore wings, traits correlate with low fly speed and acceleration, and in butterfly morphology is linked to adaptive flight behaviour remains unresolved. Here we compare the flight behaviours and wing morphologies of the spring and summer forms of two closely related butterfly species, *Pieris napi* and *P. rapae*. We first quantify three-dimensional flight behaviour by reconstructing individual flight trajectories using stereoscopic high-speed videography in an experimental outdoor cage. We then measure wing size and shape, which are characteristics assumed to influence flight behaviours in butterflies. We show that seasonal, but not interspecific, differences in flight high flight curvature. Large summer individuals exhibit rounded fore wings, which correlate with high flight speed, acceleration, turning acceleration, and advance ratio. Our study provides quantitative evidence of different flight behaviours between seasonal forms of *Pieris* butterflies, and these differences were better predicted by body size and forewing shape, rather than other factors (e.g., species identity, ambient temperature). This points toward a possible adaptive role of seasonal morphology determining flight behaviours, although this needs to be thoroughly investigated to distinguish the effects of microhabitat (e.g., behavioural thermoregulation, resource distribution between seasons, such as nectar and larval food plants), predator and parasitoid pressure, and possible sexual behavioural differences.

## Introduction

Seasonal changes in the environment induce in some species the development of polymorphic forms exhibiting traits adapted to specific environmental conditions (Xue & Leibler, 2018). Such seasonal phenotypes are typically influenced by shifts in temperature (Nijhout, 2003), species interactions, such as the availability of feeding and breeding resources (Brakefield & Reitsma, 1991) or predators’ abundance (Brakefield & Reitsma, 1991; Lyytinen et al., 2004). These adaptations are most common in regions with strong seasonality, where animals need to adjust their foraging, breeding, or dispersal strategies for a few months in the year.

For flying insects living in temperate environments, wing shape and colouration are key polyphenic traits. Polyphenism of insect wings has been shown to play roles in thermoregulation (Kingsolver, 1985), interspecific signalling (Lyytinen et al., 2004) and powered flight (Harrison, 1980). Butterflies fly mainly by using fore wings (Wootton 1992), hind wings affecting manoeuvrability (Jantzen 2008). While flight is a primary function of insect wings, little is known about its relation to wing morphology in the context of seasonal polyphenism. Differences in wing morphology are often associated with contrasted flight behaviours. For instance, wing loading (i.e. ratio of body mass to wing area) being associated with flight speed (Le Roy et al., 2019) and narrower and more elongated fore wings (higher aspect ratio) enable slow gliding and energy efficient flight, whereas more compact fore wings (lower aspect ratio) support better manoeuvrability (Le Roy et al., 2019). Less is known about how fine variations in wing shape affect flight performance (Breuker et al., 2007). Our understanding of which specific wing traits influence distinct flight behaviours, including agility and manoeuvrability, remains unclear.

Seasonal flight behaviours may be adaptive. For example, it has been hypothesised that the spring individuals of both *Pieris* species are better adapted to explore the scattered feeding and oviposition resources on their natal patch, while the summer forms might be better adapted for dispersal to novel localities (Fric et al., 2006; Shkurikhin & Oslina, 2016). Indeed, movements associated with nectaring were slower and more tortuous than movement associated with mate searching in a satyrid butterfly (Evans et al., 2020). Furthermore, increasing wing area might enhance dispersal ability, as quantified in migrating generations vs. sedentary populations of *Danaus* butterflies (Tenger-Trolander et al., 2023). Behavioural variation can also influence flight patterns in seasonal butterflies, as the preference in flight direction of migrating populations of *Pieris brassicae* seems to depend on the season of emergence (Spieth & Cordes, 2012). Thus, in the case of polyvoltine (i.e. having several generations during a year) seasonal butterflies, the factors affecting intraspecific flight behaviours likely involve both morphological and behavioural adaptations (Spieth & Cordes, 2012; Tenger-Trolander et al., 2023).

To assess whether seasonal changes in wing morphology and flight behaviours, we study two polyvoltine butterfly species, *Pieris napi* (Linnaeus, 1758) and *P. rapae* (Linnaeus, 1758) (Pieridae: Pierini). These two species are closely-related (Okamura et al., 2019), co-occur in Europe and are ecologically very similar (Tenger-Trolander et al., 2023). They are known to differ in wing size and shape in spring and summer (Fric et al., 2006; Shkurikhin & Oslina, 2016). It is thought that the spring forms of both species have a more manoeuvrable flight due to their elongated and pointed fore wings (Shkurikhin & Oslina, 2016), whereas the summer forms might have improved dispersal capacity due to their larger wings and lower wing loading (Fric et al., 2006; Shkurikhin & Oslina, 2016). Furthermore, variation in habitat use is expected among both species and seasonal forms (Friberg & Wiklund, 2019). While *P. napi* usually flies in habitats such as forest edges and feeds on biennial and perennial plants, *P. rapae* explores temporary and degraded habitats with ephemeral plants (Ohsaki & Sato, 1999), including agricultural landscapes (Ryan et al., 2019). Seasonal differences in population dynamics driven by climate have also been reported for *P. napi* and *P. rapae* (Okamura et al., 2019; von Schmalensee et al., 2023). While *P. rapae* is more abundant during summer, *P. napi* has a higher overwintering survival than *P. rapae*, resulting in higher abundances during the spring (von Schmalensee et al., 2023).

Here we quantified the flight behaviour of spring and summer forms of both *P. napi* and *P. rapae*, using a stereoscopic high-speed videography system in an outdoor cage environment. To test if seasonality drives divergent flight behaviour in *Pieris* butterflies, we compared flight characteristics (i) between *P. napi* and *P. rapae* and (ii) between seasonal forms of both species in spring and summer. More distinct differences in flight behaviour between seasonal forms than between species, correlated with certain wing morphology features, could suggest adaptive flight plasticity occurring in both *Pieris* species, which is in line with predictions that *Pieris* seasonal flight divergences are better fit to their environment (Fric et al., 2006; Shkurikhin & Oslina, 2016).

## Methods

### Butterfly flight trajectories recording

The recordings took place from the end of May to mid-August 2020. We caught *P. napi* and *P. rapae* in field during the morning (∼8:00 – 11:00) in the surroundings of České Budějovice, Czechia (48°59’28.3“N 14°26’29.6”E). *Pieris* individuals captured between May and mid-June represented the spring form, whereas the ones captured during July and August represented the summer form (von Schmalensee et al., 2023). A third generation of adults may fly during September and October during warm years (Benes et al. 2002), but these individuals have not been included in this study. The caught butterflies were stored in paper envelopes in a box with a wet napkin to avoid desiccation. Immediately after field sampling, we returned to the filming location, and flight trajectories of individual butterflies were filmed to minimalize variability in the data due to specimen handling.

Flight trajectories were filmed between 10:00 and 16:00 in an outdoor tunnel-like cage (6 m × 2.5 m × 2.5 m, length × width × height) located in the lee of a building at the Biology Centre CAS campus. The cage’s end, where we released the butterflies at the beginning of the recordings, was shaded to encourage them to fly towards the sunlit area of the cage. The individual butterflies were recorded for a maximum of 5 minutes. The tracks during which butterfly individuals fly spontaneously were included to the analyses.

Films were recorded using two GoPro Hero8 Black cameras (San Mateo, CA, USA) mounted on tripods at a height of 80 cm, and positioned at perpendicular angle to each other, following a similar setup to that of Le Roy et al., (2021). The cameras recorded at 120 frames per second using the wide lens option with a resolution of 2560×1440 pixels. To uncover within-individual variation, we filmed up to 6 flight trajectories per individual, depending on the fatigue of the individual and its ability of spontaneous flight. After the flight experiment, the butterflies were euthanized by freezing at -20°C, and later spread and mounted for morphological measurements. Ambient air temperatures during recording hours were obtained from a nearby (distance: 2 km) climatic station of the Czech Hydrometeorological Institute, České Budějovice-Rožnov.

### Quantifying flight parameters

To quantify the butterfly flight trajectories in three dimensions (3D), we used the software Argus v2.1 (Jackson et al., 2016) implemented in Python v. 3 (Van Rossum & Drake Jr., 2009). Because of the wide filming option, the coordinate systems from the camera videos were warped. We thus undistorted all videos using the Argus package DWarp. The camera setup was calibrated using the direct linear transformation (DLT) technique (Theriault et al. 2014). For this, we used a 24-cm ruler with Styrofoam balls fixed to both ends, which was moved across the cage. We then tracked the centre of the Styrofoam balls using the Argus package *Clicker*. The butterfly flight trajectories were similarly manually tracked by digitizing the positions of the butterfly thorax. But given its small size, we were not able to record the positions in certain video frames. We reconstructed the 3D coordinates per frame by merging the two synchronized 2D point trajectories and calibration coefficients using the Argus package *Wand*. The butterfly positions throughout the trajectory were then post-processed using a linear Kalman filter in MATLAB (Muijres et al., 2014) resulting in smoothed positional coordinates as well as estimated missing positions. Additionally, the total number of wingbeats per butterfly trajectory were counted during the video processing.

The flight behaviour was described using **11** parameters for every butterfly individual, all the calculations were made using a custom-written R script from Le Roy et al., (2021):

As general flight characteristics, we calculated **(1)** Wingbeat frequency (average number of wing beats per second, in Hz), **(2)** Covered distance (m), which was calculated as the consecutive distance between all tracked 3D positions, average **(3)** Flight height (m), mean **(4)** Velocity (m/s) and mean **(5)** Acceleration (m/s^2^), calculated as the first and second temporal derivative of butterfly positions through time, respectively.

As a measure of flight efficiency, we calculated mean **(6)** Advance ratio, which describes the flapping efficiency and was calculated as the ratio of mean velocity to the wingbeat frequency (Ellington, 1984).

To describe individual turning abilities, which encompass manoeuvrability and agility, we calculated **(7)** Turning acceleration, which is a component of acceleration strictly attributable to changes in flight direction, and **(8)** Turning rate, which describes how quickly the butterfly turns, which was calculated as the angular change in the direction of subsequent velocity vectors in the 3D space.

To describe the flight trajectory shape, we calculated **(9)** Sinuosity, which is the ratio of the straight distance between the starting and ending positions against the covered distance, **(10)** Curvature, which is a measure of how sharply the butterfly turns (Jantzen & Eisner, 2008), and **(11)** Ascent angle (degrees), calculated as the angle between the velocity vector and the horizontal.

### Wing morphometrics

All butterflies were mounted and photographed with a millimetre scale, using a Canon EOS250D camera with an 18-55mm EF-S lens (Tokyo, Japan). We measured conventional wing morphometrics (Figure 1) using ImageJ v. 1.53 (Schneider et al., 2012) and, for wing area calculations, we used GIMP (GIMP Development Team, 2019). We measured the left wings for all calculations; if the left wings were damaged, the right wings were used. For all specimens, we calculated Fore wing length (cm), which is the distance between the wing base (landmark point 1; Figure 1) and the outer edge of vein 13 (Bai et al. 2015, Figure 1); Fore wing width (cm), which is the highest width perpendicular to the measured fore wing length; Aspect ratio, which characterises the relative elongation and narrowness of the forewings and is calculated by the ratio between fore wing length and width; Fore wing area (cm^2^); Thoracic volume (cm^3^), which served as proxy to body mass, and was calculated as the cylindrical volume inferred from the thorax length and width; Total wing area (cm^2^), measured as twice the fore wing area (dorsal view) plus twice the hindwing area (ventral view), using the histogram function of GIMP; and Wing loading, being the ratio of thoracic volume (as proxy for mass) and the total wing area.

**Figure 1.**
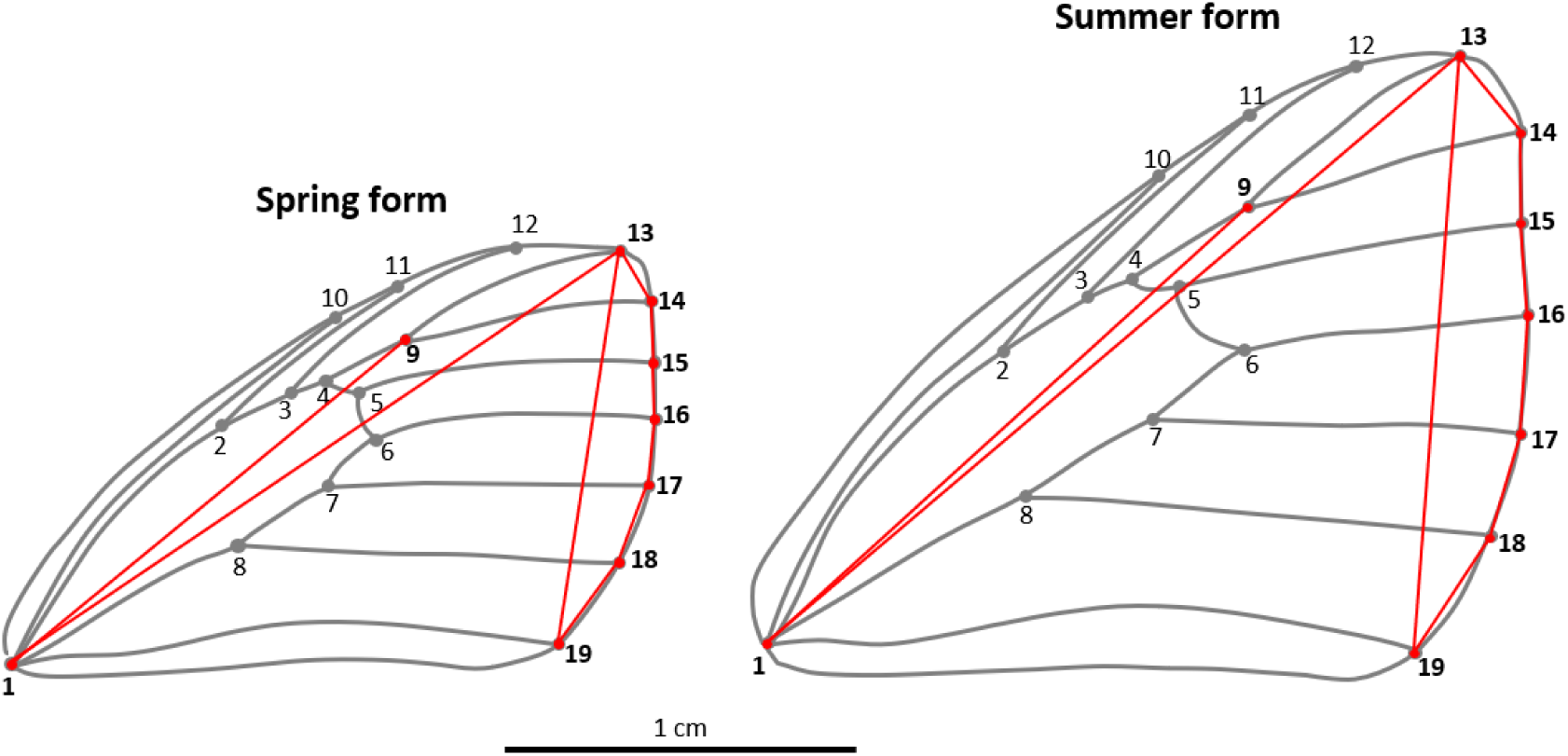
Wing morphology of *Pieris* white butterflies and their seasonal forms. The two indices of wing shape were calculated: Relative length of the marginal region (the distance between landmark 1 to 9 divided by the straight distance between landmarks 1 to 13); Curvature of the fore wing outer edge (the sum of distances around the outer edge from landmarks 13 to 19 divided by straight distance between landmarks 13 and 19).

In addition, we calculated two wing shape characteristics, which we hypothesise to affect flight. These wing shape characteristics differ between species and between seasonal forms based on description of Shkurikhin & Oslina, 2016, who quantified morphological difference of seasonal forms of *P. napi* and *P. rapae*. **-**First, Curvature of the fore wing outer edge, which is known to differ between the spring and the summer forms (Shkurikhin & Oslina, 2016), was calculated as the sum of distances around the terminal veins at the outer edge (landmark points 13, 14, 15, 16, 17, 18 and 19, respectively) divided by the straight distance between veins landmark points 13 to 19 (Figure 1). Curvature of the fore wing outer edge close to 1 represents the rounded forewings of summer *Pieris*, while an index higher than 1 depicts the elongated fore wings of spring *Pieris* (Shkurikhin & Oslina, 2016). Then, we calculated the Relative length of the marginal region, measured as the distance between landmark 1 to 9 divided by the straight distance between landmarks 1 to 13 ((Figure 1). Relative length of the marginal region differs among the species based on Shkurikhin & Oslina, 2016. The shape characteristics of one *P. napi* (specimen b220) were not possible to calculate as its wing margins were damaged during handling after the video recordings; the specimen was excluded from the analyses of the effect of wing shape on flight behaviour. We use *aov* function in the R v. 4.2.1 (R Core Team, 2022) to test whether morphological variables potentially affecting flight (Fore wing area, Aspect ratio, Wing loading, Curvature of the fore wing outer edge and Relative length of the marginal region) differ among species, seasonal forms and sexes, including interactions between these factors.

### Interspecific and seasonal differences in flight behaviour

We compared differences in flight behaviour between the species (*P. napi* and *P. rapae*), and between their seasonal forms, spring (May to mid-June) and summer (July and August). First, the measured flight characteristics were reduced in dimensionality and visualized using Principal Component Analysis (PCA) in the R software. The function ‘*imputePCA*’of the package *missMDA* (Josse & Husson, 2016) was used to estimate missing values of average wingbeat frequency and advance ratio in four 3D trajectories where wingbeats were difficult to score. To visualize possible interspecific and seasonal differentiation in flight behaviour, we used the function *fviz_pca* of the package *devtools* (Wickham et al., 2022).

Second, the differences in flight behaviour among species and their seasonal forms were estimated using linear models with mixed effects (LMMs), as implemented in the R package *lme4* (Bates et. al. 2015) via the function ‘*lmer*’. The response variables were log/transformed flight characteristics (1-11), except of (2) Covered distance, fitted in separate models with species, seasonal form and sex identity as predictor variables. The log-transformed (2) Covered distance was included as a fixed factor, because it was used to derive other flight characteristics describing trajectories or flight dynamics. As we recorded multiple trajectories per individual, specimen identity was included as a random factor. Then, post hoc, pairwise comparisons were performed for flight characteristics which were found to differ significantly among species, seasonal forms or sexes in the previous LMM analyses. We used Tukey post hoc tests implemented in the R package *emmeans* (Lenth, 2022) to calculate mean values of the flight characteristics for each species and seasonal form and estimated marginal (*emmeans*) values of the LMMs. We tested the effect of ambient temperature on flight characteristics in LMM with species and seasonal form as fixed factor and individual as random factor and then in simplified LMM with ambient temperature as explaining variable and individual as random factor.

### Associations of wing morphologies with flight behaviour

We additionally tested the effect of morphology on flight while excluding species and seasonal form to focus on how morphology affects flight. To test whether wing morphological traits of recorded individuals were associated with specific flight characteristics, we used LMMs with flight characteristics as response variables in separate models. As explanatory variables we used log transformed Fore wing area, Aspect ratio, Wing loading, Curvature of the fore wing outer edge and Relative length of the marginal region. The log-transformed (2) Covered distance and sex were included as fixed effects and specimen identity as a random factor. The effect of morphology was tested without including species and seasonal form as fixed factor, to reveal solely effect of morphology on flight.

## Results

### Interspecific and seasonal differences in morphology

We quantified 106 flight trajectories of 31 butterfly individuals: 14 *P. rapae* and 17 *P. napi* (Table 1). The differences in wing morphology between the two *Pieris* species were lower than between their seasonal forms. The two *Pieris* species differed only in Relative length of marginal region (F_1,23_ = 9.111, p =0.006), as *P. napi* had a longer Relative length of marginal region than *P. rapae* (Figure 1, Table 1). *Pieris napi* and *P. rapae* did not significantly differ in their Fore wing area (F_1,24_ = 0.0004, p = 0.863), Aspect ratio (F_1,24_=0.007, p = 0.936), Wing loadings (F_1,24_ = 0.130, p = 0.722) and Curvature of the fore wing outer edge (F_1,24_=1.037, p = 0.319).

**Table 1.** Overview table summarising the number of male and female individuals of *Pieris napi* and *P. rapae* butterflies and their seasonal forms, number of tracks recorded, and mean values (±SE) of Fore wing area, Aspect ratio (the ratio of length to the width of wing), Wing loading (the ratio of Thoracic volume and Total wing area), Curvature of the fore wing outer edge and Relative length of marginal region (see Figure 1). *Pieris* species differed only in Relative length of marginal region, however, their seasonal forms differed in multiple morphological variables. Generally, the spring form is smaller with elongated fore wings. The summer form was larger with more rounded fore wings.

|  | N | Spring<br>/Summer | Nu of<br>tracks | Fore wing<br>area<br>(mm<sup>2</sup>) | Wing<br>loading<br>(mm<sup>3</sup>/mm<sup>2</sup>) | Aspect<br>ratio | Outer<br>edge | Marginal<br>region |
| --- | --- | --- | --- | --- | --- | --- | --- | --- |
| **Males** |  |  |  |  |  |  |  |  |
| *napi* | 14 | 3/11 | 51 | 248<br>(8.93) | 0.0223<br>(0.00182) | 1.75<br>(0.015) | 1.09<br>(0.008) | 0.72<br>(0.008) |
| *rapae* | 12 | 5/7 | 38 | 242<br>(9.54) | 0.0214<br>(0.00203) | 1.74<br>(0.02) | 1.09<br>(0.009) | 0.68<br>(0.007) |
| Spring | 8 |  | 22 | 225.6<br>(8.29) | 0.0249<br>(0.00242) | 1.77<br>(0.02) | 1.11<br>(0.012) | 0.70<br>(0.013) |
| Summer | 18 |  | 67 | 256.3<br>(7.93) | 0.0209<br>(0.00156) | 1.73<br>(0.01) | 1.080<br>(0.006) | 0.71<br>(0.007) |
| **Females** |  |  |  |  |  |  |  |  |
| *napi* | 3 | 1/2 | 12 | 212<br>(29.1) | 0.0164<br>(0.0022) | 1.74<br>(0.055) | 1.10<br>(0.020) | 0.70<br>(0.010) |
| *rapae* | 2 | 0/2 | 5 | 243<br>(22.4) | 0.0160<br>(0.00104) | 1.80<br>(0.015) | 1.06<br>(0.012) | 0.69<br>(0.033) |
| Spring | 1 |  | 4 | 154<br>(NA) | 0.0021<br>(NA) | 1.85<br>(NA) | 1.135<br>(NA) | 0.70<br>(NA) |
| Summer | 4 |  | 13 | 242<br>(9.2) | 0.0015<br>(0.00008) | 1.74<br>(0.035) | 1.07<br>(0.006) | 0.69<br>(0.016) |

Seasonal forms differed in size and shape of fore wings (Table 1). For both species, spring individuals were smaller than summer ones. Spring *Pieris* had significantly lower Fore wing area (F_1,24_**=** 9.934, p = 0.004) than summer *Pieris*. Although only marginally significant, Wing loading (F_1,24_ **=** 3.447, p = 0.076) and Aspect ratio (F_1,24_ **=**3.956, p = 0.058) tended to be higher in the spring forms (Table 1). Higher values of Aspect ratio describe long and slender wings, the higher Wing loading describe lower wing area in relation to given body mass. Curvature of the fore wing outer edge was significantly higher in spring *Pieris* (F_1,23_ = 9.766, p = 0.005), meaning the spring generation had more elongated fore wings. The curvature of the fore wing outer edge was closer to 1 in summer *Pieris* meaning that the summer form had more rounded fore wings (Figure 1). Relative length of the fore wing marginal region (F_1,23_ **=** 0.045, p = 0.834) did not differ between spring and summer forms. We have not detected significant differences in morphological traits between sexes.

### Interspecific and seasonal differences in flight

The first axis of the PCA (32.8% variance explained) was positively associated with higher (4) Velocity, (5) Acceleration, (6) Advance ratio, and (7) Turning acceleration, and lower (3) Flight height and (10) Curvature. The second PCA axis (20.63% explained variance) was positively associated with (11) Ascent angle and negatively with mean (8) Turning rate and (9) Sinuosity. The PCA space did not reveal substantial differences between individuals of *P. napi* and *P. rapae*, which had a large variation in the measured flight parameters (Figure 2). Conversely, the flight behaviours of the spring and summer forms clustered separately, although higher variability in flight characteristics was present within summer individuals. Spring individuals had lower (6) Advance ratio, (4) Velocity, (5) Acceleration and (7) Turning acceleration, but higher flight (10) Curvature and (3) Flight height compared to the summer form.

**Figure 2.**
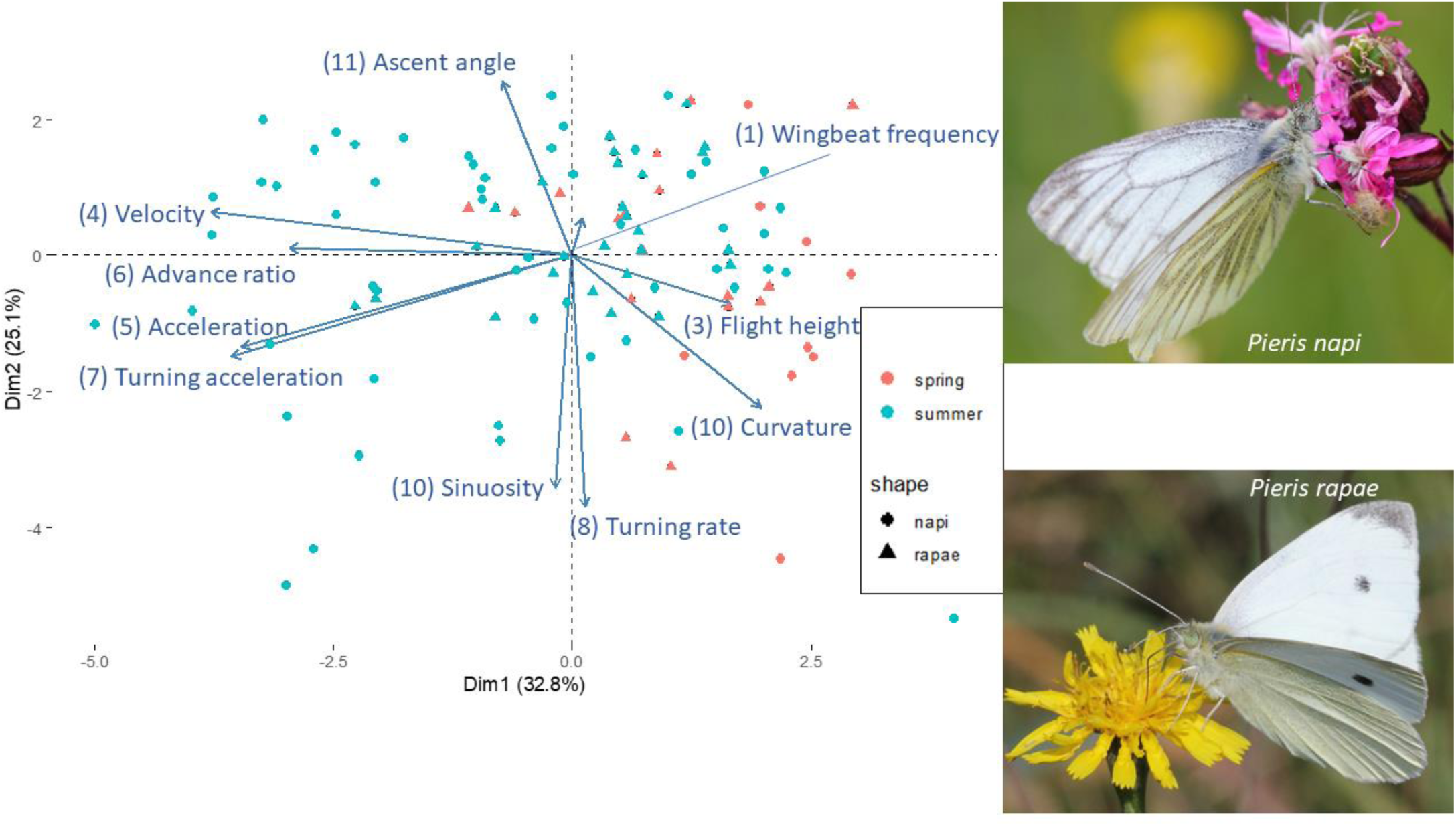
Seasonal forms but not species of *Pieris* butterflies differed in their flight characteristics. The first and second axes of PCA analyses of flight characteristics of the two closely related sympatric *P. napi* and *P. rapae* (Pieridae) and their seasonal, spring and summer, forms. Each point represents a recorded trajectory (photo by Zdeněk Hanč).

Based on the results of the LMMs, the two *Pieris* species did not significantly differ in any of the measured flight characteristics (Table 2). Further, males and females did not differ significantly in their tested flight characteristics, which might be caused by the low number of recorded females (Table 1). However, males tend to have higher trajectory (10) Curvature than females (F=3.413, df = 30.06, p = 0.075); the mean curvature recorded in males was 3.15 (±0.520) and in females 1.85 (±0.416). In contrast, the seasonal forms differed significantly (Table 2), in agreement with the clustering pattern highlighted by the PCA, spring individuals had lower (6) Advance ratio (p < 0.01), (4) Velocity (p < 0.01), (5) Acceleration (p = 0.03), (7) Turning acceleration (p < 0.01) and (8) Turning rate (p < 0.01), but higher flight (10) Curvature (p < 0.01) than summer individuals. Range of the air temperature during recording hours was 14°C to 36 °C, mean value was19(±0.3) °C. We did not detect any significant effect of the air temperature during recording hours on flight behaviour.

**Table 2.** In an outdoor cage experiment *Pieris napi* and *P. rapae* butterfly species did not differ in flight behaviour, but their seasonal forms do. Comparison of interspecific differences in flight characteristics in two closely related species, *P. napi* (17 individuals, 63 trajectories) and *P. rapae* (14 individuals, 43 trajectories), and between their seasonal forms, spring (9 individuals, 26 trajectories) and summer (22 individuals, 80 trajectories). For each *Pieris* individual, flight characteristics were inferred for several trajectories of differing length, thus linear mixed models (LMMs) include individual identity as a random factor and log-transformed (2) Covered distance and sex as fixed factor. For flight characteristics, which significantly differed among seasonal forms were estimated marginal means of LMM (emmean) transformed to real values of the flight characteristics. Significant relationships are highlighted **in bold**.

| Flight characteristics | Species effect |  |  | Seasonal form effect |  |  |  |  |
| --- | --- | --- | --- | --- | --- | --- | --- | --- |
|  | F | Dendf | P | F | Dendf | P | emmean (SE) |  |
|  |  |  |  |  |  |  | spring<br>(N=26) | summer<br>(N=80) |
| (1) Wingbeat frequency | 0.393 | 30.111 | 0.54 | 0.532 | 32.548 | 0.47 |  |  |
| (3) Flight height | 0.001 | 28.692 | 0.97 | 30.283 | 30.283 | 0.34 |  |  |
| **(4) Velocity** | 0.006 | 27.087 | 0.94 | **10.928** | **28.864** | **<0.01** | **1.2**<br>**(0.124)** | **1.7**<br>**(0.118)** |
| **(5) Acceleration** | 0.008 | 25.561 | 0.93 | **5.001** | **27.238** | **0.03** | **2.31**<br>**(0.400)** | **3.43**<br>**(0.398)** |
| **(6) Advance ratio** | 8.651 | 27.886 | 0.67 | **8.651** | **30.614** | **<0.01** | **0.10**<br>**(0.013)** | **0.14**<br>**(0.012)** |
| **(7) Turning acceleration** | 0.046 | 25.746 | 0.83 | **10.438** | **27.538** | **<0.01** | **1.30**<br>**(0.232)** | **2.34**<br>**(0.278)** |
| (8) Turning rate | 0.002 | 28.751 | 0.96 | 0.115 | 30.766 | <0.01 |  |  |
| (9) Sinuosity | 0.260 | 33.226 | 0.26 | 0.762 | 35.440 | 0.389 |  |  |
| **(10) Curvature** | 0.014 | 31.372 | 0.91 | **13.46** | **33.501** | **<0.01** | **3.80**<br>**(0.927)** | **1.53**<br>**(0.249)** |
| (11) Ascent angle | 0.334 | 28.215 | 0.57 | 0.911 | 30.494 | 0.35 |  |  |

### Associations of wing morphologies with flight behaviour

Flight characteristics were affected by Fore wing area, Aspect ratio and Curvature of the fore wing outer edge (Table 3), but not by Wing loading and Relative length of marginal region. Curvature of the fore wing outer edge was the most significant morphological variable correlated with different flight behaviour of the two seasonal forms (Table 3). The higher Curvature of the fore wing outer edge (i.e., typical for the elongated fore wings found in spring *Pieris*) were associated with higher flight (10) Curvature (p = 0.01), whereas the lower values of Curvature of the fore wing outer edge (i.e., the rounded fore wings in summer *Pieris*) were associated with higher (4) Velocity (p = 0.03) and higher (6) Advance ratio (p = 0.05). The larger Fore wing area, found in summer *Pieris*, positively correlate with higher (4) Velocity (p = 0.04), (6) Advance ratio (p = 0.02) and tended to also correlate with higher (5) Acceleration (p = 0.06) (Table 3). Butterflies with lower Aspect ratio (i.e. more rounded wings of the summer form) tend to have higher (6) Advance ratio (p = 0.07), this result corresponds with the higher (6) Advance ratio recorded for the summer form (Table 2). Although we have detected a significantly higher (7) Turning acceleration (measure of turning ability; p < 0.01) in summer *Pieris* (Table 2), we did not find any measured wing morphology trait connected to such a difference. Contrary to previous assumptions, we have not detected any significant effect of Wing loading on the measured flight parameters between seasonal forms in *P. napi* and *P. rapae*. This might be caused by using Thoracic volume to infer Wing loading, instead of fresh weights.

**Table 3.**
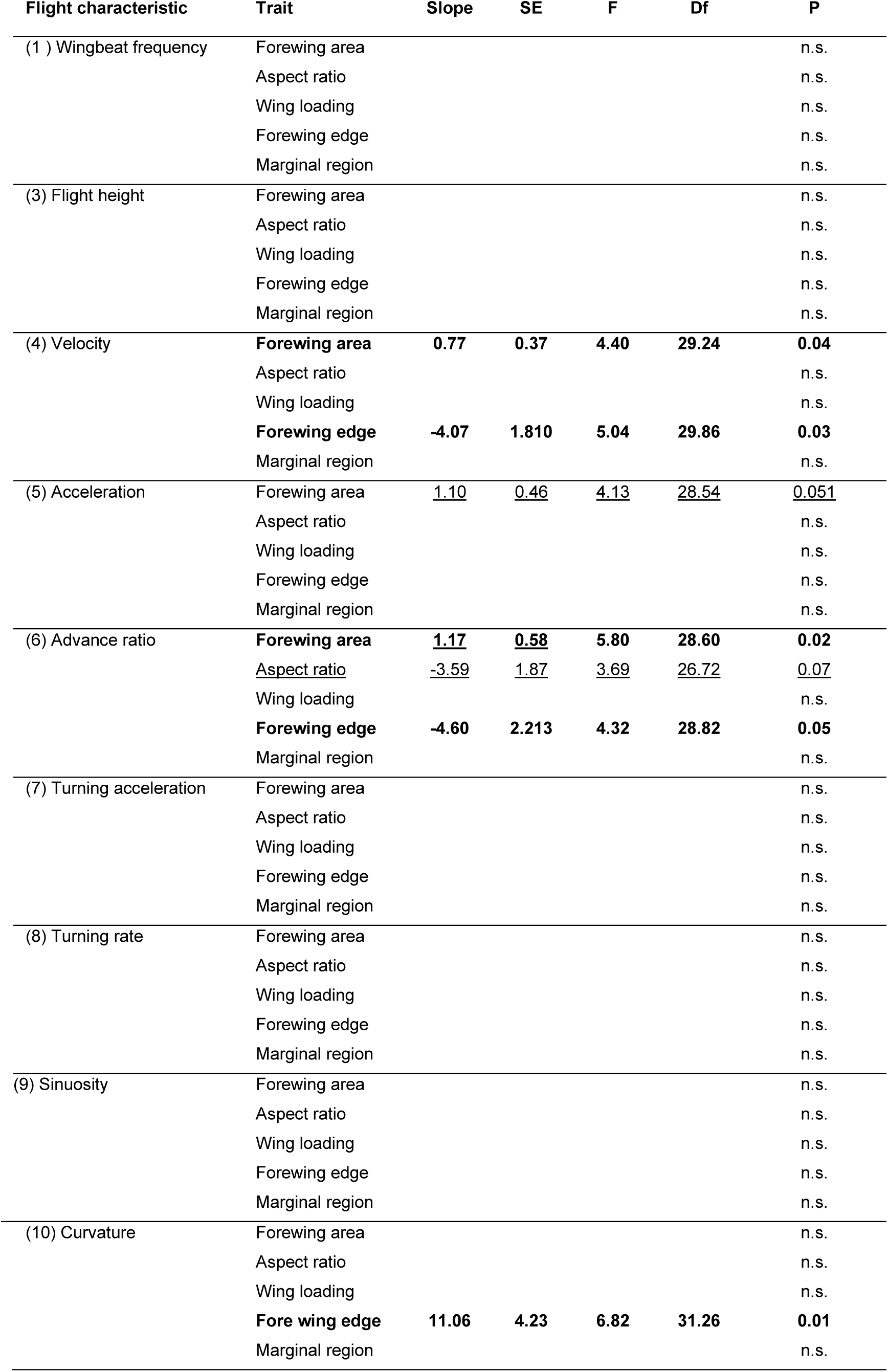
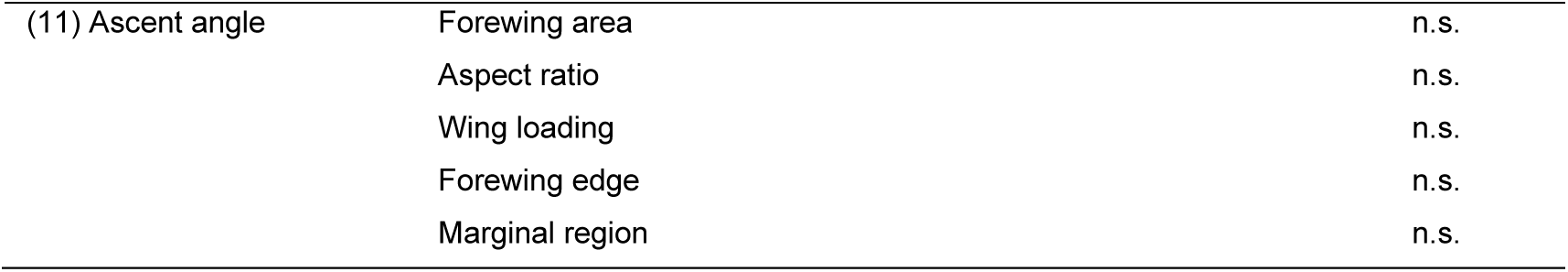
Flight performance of *Pieris napi* and *P. rapae* (Pieridae) butterflies was determined by Fore wing area, Aspect ratio and Curvature of the fore wing outer edge (Forewing edge) , but not by Wing loading and Relative length of marginal region (Marginal region).

| Flight characteristic | Trait | Slope | SE | F | Df | P |
| --- | --- | --- | --- | --- | --- | --- |
| (1 ) Wingbeat frequency | Forewing area |  |  |  |  | n.s. |
|  | Aspect ratio |  |  |  |  | n.s. |
|  | Wing loading |  |  |  |  | n.s. |
|  | Forewing edge |  |  |  |  | n.s. |
|  | Marginal region |  |  |  |  | n.s. |
| (3) Flight height | Forewing area |  |  |  |  | n.s. |
|  | Aspect ratio |  |  |  |  | n.s. |
|  | Wing loading |  |  |  |  | n.s. |
|  | Forewing edge |  |  |  |  | n.s. |
|  | Marginal region |  |  |  |  | n.s. |
| (4) Velocity | **Forewing area** | **0.77** | **0.37** | **4.40** | **29.24** | **0.04** |
|  | Aspect ratio |  |  |  |  | n.s. |
|  | Wing loading |  |  |  |  | n.s. |
|  | **Forewing edge** | **-4.07** | **1.810** | **5.04** | **29.86** | **0.03** |
|  | Marginal region |  |  |  |  | n.s. |
| (5) Acceleration | Forewing area | <u>1.10</u> | <u>0.46</u> | <u>4.13</u> | <u>28.54</u> | <u>0.051</u> |
|  | Aspect ratio |  |  |  |  | n.s. |
|  | Wing loading |  |  |  |  | n.s. |
|  | Forewing edge |  |  |  |  | n.s. |
|  | Marginal region |  |  |  |  | n.s. |
| (6) Advance ratio | **Forewing area** | **<u>1.17</u>** | **<u>0.58</u>** | **5.80** | **28.60** | **0.02** |
|  | <u>Aspect ratio</u> | <u>-3.59</u> | <u>1.87</u> | <u>3.69</u> | <u>26.72</u> | <u>0.07</u> |
|  | Wing loading |  |  |  |  | n.s. |
|  | **Forewing edge** | **-4.60** | **2.213** | **4.32** | **28.82** | **0.05** |
|  | Marginal region |  |  |  |  | n.s. |
| (7) Turning acceleration | Forewing area |  |  |  |  | n.s. |
|  | Aspect ratio |  |  |  |  | n.s. |
|  | Wing loading |  |  |  |  | n.s. |
|  | Forewing edge |  |  |  |  | n.s. |
|  | Marginal region |  |  |  |  | n.s. |
| (8) Turning rate | Forewing area |  |  |  |  | n.s. |
|  | Aspect ratio |  |  |  |  | n.s. |
|  | Wing loading |  |  |  |  | n.s. |
|  | Forewing edge |  |  |  |  | n.s. |
|  | Marginal region |  |  |  |  | n.s. |
| (9) Sinuosity | Forewing area |  |  |  |  | n.s. |
|  | Aspect ratio |  |  |  |  | n.s. |
|  | Wing loading |  |  |  |  | n.s. |
|  | Forewing edge |  |  |  |  | n.s. |
|  | Marginal region |  |  |  |  | n.s. |
| (10) Curvature | Forewing area |  |  |  |  | n.s. |
|  | Aspect ratio |  |  |  |  | n.s. |
|  | Wing loading |  |  |  |  | n.s. |
|  | **Fore wing edge** | **11.06** | **4.23** | **6.82** | **31.26** | **0.01** |
|  | Marginal region |  |  |  |  | n.s. |

**Table 3.** Flight performance of *Pieris napi* and *P. rapae* (Pieridae) butterflies was determined by 713 Fore wing area, Aspect ratio and Curvature of the fore wing outer edge (Forewing edge) , but not 714 by Wing loading and Relative length of marginal region (Marginal region).
|  |  |  |
| --- | --- | --- |
| (11) Ascent angle | Forewing area | n.s. |
|  | Aspect ratio | n.s. |
|  | Wing loading | n.s. |
|  | Forewing edge | n.s. |
|  | Marginal region | n.s. |

## Discussion

In this study, we used stereoscopic high-speed videography to quantify and compare the flight behaviours of two closely related *Pieris* species and their seasonal forms in an outdoor cage experiment. We hypothesise, that *P. napi* associate with forest edges may fly with higher sinuosity and manoeuvrability compared to straighter flight in *P. rapae*, which is associated with agricultural open landscapes. However, the hypothesised divergence in flight behaviours between species due to different habitat preferences was not supported by our findings, suggesting that flight adaptation to slightly different habitats use has not evolved in the studied species. Interestingly, our results suggest that seasonal variation in wing size and shape is associated with contrasting flight characteristics (Table 2). The morphological differences among seasonal forms are thought to be adaptative, with spring form being more sedentary and the summer form more dispersive (Fric et al., 2006, Shkurikhin & Oslina, 2016). On one hand, the elongated fore wings of smaller spring *Pieris* (Fric et al., 2006; Shkurikhin & Oslina, 2016) are associated with lower velocity, acceleration, advance ratio and turning acceleration, and higher flight curvature, indicating manoeuvrable flight with tortuous trajectories. On the other hand, larger and rounded fore wings of summer *Pieris* are associated with higher velocity, acceleration, advance ratio and turning acceleration, but lower curvature. By recording the flight of wild-caught individuals in outdoor cages exposed to seasonal variation of ambient temperatures, we expect that the recorded flight behaviour would mirror those displayed by butterflies in natural conditions. Although our experiment provides a first insight into the differentiation of flight behaviour between seasonal forms, experiments with captive bred individuals in controlled conditions are needed to ascertain whether the differences between seasonal forms might be adaptive.

Differentiation of adult morphology (size and shape) and, thus, possibly also indirectly flight behaviour, are determined by larval development. Larval growth is affected by host plant species and their quality (Hwang et al., 2008), by ambient temperatures and photoperiod (Nylin, 1994). Low ambient temperatures experienced by immatures lead to the development of larger adult individuals of summer form, which may indirectly influence their faster flight compared to smaller individuals reared in colder conditions (Büyükyilmaz & Tseng, 2022). Shortening days towards the end of summer and autumn stimulate the initiation of the overwintering stage, pupae, resulting in smaller spring individuals (Wiklund et al. 1991). In addition, wing shape differentiation is driven by hormonal pathways influenced by an array of environmental factors (Nylin, 1994, Bai et al., 2015). We propose that wing shape may be adaptive and influenced by selection for flight performance, as suggested by Fric et al. (2006) and Shkurikhin & Oslina (2016), whereas differences in wing size and area may be determined by larval diet and ambient temperatures. Nonetheless, regardless of whether seasonal flight behaviour differences in *Pieris* are adaptive or plastic, their flight behaviours would likely be affected by larval developmental disruptions driven by ongoing climatic changes.

Our approach revealed significant differences in flight behaviour between seasonal forms of *Pieris* and the effects of wing morphology on flight. However, using wild specimens for our study likely resulted in uncontrolled variables such as variation in age, mating or feeding status, possibly impacting flight behaviour (Almbro & Kullberg, 2008, 2012). Similarly, we have not revealed the effect of sex (Wiklund et al. 1991, Van Dyck & Wiklund, 2002), as only 5 females were collected (Table 1). For example, sexual selection favouring larger size in males can differ between seasons in *P. napi* (Wiklund et al. 1991). Further studies should investigate the effects of varying environmental conditions (air temperature, host plant, etc.) during larval and adult stages on flight behaviour and performance.

### Seasonal but not interspecific morphological differences affect flight behaviour

Differences in environmental conditions between spring and summer may have driven divergent evolution of flight behaviours in *Pieris* butterflies (Table 2). While the two closely related species *P. napi* and *P. rapae* have variable, but overall undistinguishable flights and morphologies (Table 2), their respective seasonal forms appear to diverge in morphology (Table 1) and flight behaviour (Table 2). The observed morphological differences between seasonal forms are consistent with previous studies. Fric et al. (2006) and Shkurikhin & Oslina (2016) reported that spring individuals were smaller with more elongated wings and higher wing loading compared to larger summer individuals with more rounded wings. They hypothesised that the spring individuals explore resources on their natal patch, while the summer individuals disperse to novel localities (Fric et al., 2006; Shkurikhin & Oslina, 2016). Dispersal may indeed be favoured because it permits access to more newly emerged larval feeding plants and possibly reduced parasitoid pressure (Kerr et al., 2020; Ohsaki & Sato, 1999).

Based on the morphological differences, it was generally assumed that spring individuals, with their high wing loading, fly faster and with greater manoeuvrability than the summer individuals (Fric et al., 2006; Shkurikhin & Oslina, 2016). Contrary to such expectations, the flight of spring *Pieris* was significantly slower than that of summer individuals (Table 2). A lower flight speed in smaller individuals of *P. rapae* – albeit showing higher wing loading – was also observed by Büyükyilmaz & Tseng (2022), who used a flight mill to measure velocity and covered distance of those butterflies. Similarly, Almbro and Kullberg (2012) noted that larger wing loading is associated with reduced flight speed in *P. napi* males, but not in females. Altogether, the available quantitative evidence suggests that larger wing size, rather than higher wing loading, may result in higher flight speed. Alternatively, these differences in flight behaviour may not be solely conditioned by morphology. Indeed, determining the effect of morphology on performance is often challenging, as differences may only be apparent when individuals are pushed to their limits (Losos et al., 2002). Despite the general expectation that flight speed increases with wing loading (Le Roy et al., 2019), *Pieris* contradicts this trend, with spring forms having higher wing loading (Table 1; Fric et al., 2006) yet significantly lower flight velocities than summer *Pieris*.

Our experimental setup did not allow for a direct assessment of the effect of temperature on flight behaviour. The lower flight velocity observed in spring *Pieris* may be attributed to lower ambient temperature, which in turn lower body temperatures and consequently affect flight performance (Tsuji et al., 1986). Further, it was expected that flight duration would be constrained by the cooler spring temperatures, leading butterflies to land more frequently to warm up in sunlight (Shkurikhin & Oslina, 2016). However, we did not reveal any significant effects of ambient air temperature on flight behaviour using data from a nearby weather station. Flight speed has also been reported to be primarily determined by morphology (slow flight predicted by small body size in *P. rapae*), rather than by ambient temperature, in an experiment conducted under controlled climatic conditions (Büyükyilmaz & Tseng, 2022). Thus, the available evidence suggests that ambient temperature is probably not the primary direct factor driving seasonal differences in *Pieris* flight behaviour.

Spring form was predicted to have more manoeuvrable flights compared to the straight flight of the dispersive summer form (Fric et al., 2006; Shkurikhin & Oslina, 2016). Indeed, we provide quantitative evidence for higher manoeuvrability in spring individuals, suggested by a higher flight curvature (Table 2), which was significantly associated with the fore wing shape. The small and elongated fore wings typical for the spring individuals were linked to flight trajectories with higher curvature (Table 3). Such flight characteristics might increase exploration efficiency in individuals living during early spring (Root & Kareiva, 1984), which may be beneficial for locating emerging nectar sources and larval feeding plants (Danks, 2007). Although the benefits of adopting a curvy flight pattern for foraging in a dispersed resource environment remains hypothetical, the higher flight curvature of spring *Pieris* may have evolved jointly with a specialised foraging strategy. Next, males and females may exhibit distinct flight patterns across the landscape due to differences in their resource utilization strategies and habitat use (Evans et al. 2020). In that sense, the divergent flight behaviour of spring *Pieris* might be adaptive and linked to movements across the landscape, which might result in more effective exploitation of newly emerging feeding and oviposition resources (Cant et al., 2005).

The summer dispersal of *Pieris* was hypothesised to facilitate the colonization of novel patches with larval feeding plants and reduces parasitoid pressure on the novel sites (Kerr et al., 2020; Ohsaki & Sato, 1999). The dispersal was hypothesised to be supported by the larger wing area and, consequently, the lower wing loading typical of summer *Pieris* (Fric et al., 2006; Shkurikhin & Oslina, 2016). But it was also postulated, that more rounded wings, might be adaptive for using ascending thermal air currents for dispersal (Fric et al., 2006). Indeed, it was the rounded fore wing shape of the summer form that was significantly associated with high advance ratio, thus, might contribute to effective dispersal (Table 3). Next, we found that larger wing area, but not wing loading, seems to be positively correlated with advance ratio and acceleration, though with marginal statistical significance (Table 3). Furthermore, we did not find evidence supporting that the larger wing size of summer *Pieris* decreases acceleration predicted by Shkurikhin & Oslina (2016). On the contrary, we found higher acceleration and turning acceleration in larger individuals and the summer form of *Pieris*. Such a powered flight may reflect the fast linear flight used during dispersal (Cant et al., 2005), but it is also recognised that dispersal flight is affected by warmer summer weather (Cormont et al., 2011). The higher acceleration and turning acceleration of the summer form might also represent seasonal behavioural response to increased predator pressure on adult butterflies during summer for example by naïve juvenile birds (Zvereva & Kozlov, 2021) or dragonflies (Sang & Teder, 2011). This hypothesis should however be taken with caution because it has also been predicted that the predation pressure on insects is likely higher during spring than during summer in Europe (e.g., Scandinavia, Remmel et al., 2009) and probably also in the Czech Republic. Although *Pieris* butterflies are palatable, free-flying *Pieris* were also reported to be actively avoided by birds (Lyytinen et al., 1999). Thus, we argue that the faster flight with higher accelerations recorded in summer forms could also represent an adaptive response in relation to predation.

Our results highlight the high variability in butterfly flight behaviours, which challenge the constancy of simple biomechanics assumptions based on wing morphology alone (for a review, see Le Roy et al., 2019). The main morphological trait that might be involved in the flight divergence observed between spring and summer *Pieris* is the shape of the fore wings (Figure 1, Table 3). Wing shape variation in temperate butterflies might be either plastic or have a genetic background, as revealed in *Pararge aegeria* (Nymphalidae) (Van Dyck & Wiklund, 2002, Berwaerts & Van Dyck, 2004). Similarly, subtle differences in wing shape are linked with distinct dispersal capacity in females of *Melitaea cinxia* (Nymphalidae) (Breuker et al., 2007). We also cannot rule out that the seasonal wing morphologies may be related to differences in thermoregulation properties, in addition to divergent flight characteristics. Indeed, insect wing is a complex trait, evolving under the effect multiple and sometimes conflictive selective pressures (Wootton 1992). The differences in shape could be driven by developmental conditions (Büyükyilmaz & Tseng, 2022) and differ among sexes (Breuker et al., 2007). Thus, the morphological divergence between seasonal forms may be conditioned by multiple selective pressures. Our results add to mounting evidence that the predictors of flight behaviour such as manoeuvrability and dispersal capacity are complex and are likely the result of both morphological adaptation and behavioural variation.

## Conclusions

Our study reveals different flight behaviours between seasonal forms of two *Pieris* species. We provide quantitative evidence relating wing morphology, in particular the shape of fore wings and fore wing area, to differences in flight behaviour (Le Roy et al., 2019). Different foraging flight behaviours (e.g., higher flight curvature of spring *Pieris* in relation to resource exploration) possibly promote the seasonal divergence in *Pieris* flight, which has rarely been documented in butterflies (e.g., Dell’Aglio et al., 2022; Mena et al., 2020; Spieth & Cordes, 2012). In the future, a combination of experiments with laboratory bred individuals (Almbro & Kullberg, 2008) together with recording the flight behaviour of wild butterflies in both, controlled and natural settings (e.g., Cant et al., 2005; Maggiora et al., 2019) will improve our understanding of the mechanisms driving seasonal differences in flight behaviour.

## Acknowledgements

We would like to extend our gratitude towards Alena Sucháčková Bartoňová and Klára Aurová to help with sampling of adult butterflies as well as the student assistants Klára Hájková who helped with the time-consuming task of video tracking and Zdeněk Hanč for providing photos of the species for the graphical abstract.

## Authors contributions

**I.K.** - Conceptualization, Investigation, methodology, data curation and analysis, writing original draft, review and editing. **D.L.** - data analysis, writing original draft, review and editing. **F.M.R.** - Conceptualization, Investigation, and editing, **L.R. –** data curation and investigation, review and editing, **C.L.R.** - data analysis, review and editing, **P.M.M.** - Funding acquisition, investigation, methodology, conceptualization, review and editing.

## Data Accessibility

All data and scripts are archived in the Figshare repository, (DOI: https://doi.org/10.6084/m9.figshare.24631410.v4).

## References

Almbro, M., & Kullberg, C. (2008). Impaired escape flight ability in butterflies due to low flight muscle ratio prior to hibernation. Journal of Experimental Biology, 211(1), 24–28. 10.1242/jeb.008219

Almbro, M., & Kullberg, C. (2012). Weight loading and reproductive status affect the flight performance of *Pieris napi* butterflies. Journal of Insect Behavior, 25(5), 441–452. 10.1007/s10905-011-9309-1

Bai, Y., Ma, L. B., Xu, S. Q., & Wang, G. H. (2015). A geometric morphometric study of the wing shapes of *Pieris rapae* (Lepidoptera: Pieridae) from the Qinling Mountains and adjacent regions: An environmental and distance-based consideration. Florida Entomologist, 98(1), 162–169.

Benes, J. Konvicka, M., Dvorak, J., Fric, Z., Havelda, Z., Pavlicko, A., Vrabec, V. & Weidenhoffer, Z. (eds) (2002) Butterflies of the Czech Republic: Distribution and Conservation I., II. SOM, Prague, Czech Republic.

Berwaerts, K., & Van Dyck, H. (2004). Take-off performance under optimal and suboptimal thermal conditions in the butterfly *Pararge aegeria*. Oecologia, 141(3), 536–545. 10.1007/s00442-004-1661-9

Berwaerts, K., Van Dyck, H., & Aerts, P. (2002). Does flight morphology relate to flight performance? An experimental test with the butterfly *Pararge aegeria*. Functional Ecology, 16(4), 484–491. 10.1046/j.1365-2435.2002.00650.x

Bladon, A. J., Lewis, M., Bladon, E. K., Buckton, S. J., Corbett, S., Ewing, S. R., Hayes, M. P., Hitchcock, G. E., Knock, R., Lucas, C., McVeigh, A., Menéndez, R., Walker, J. M., Fayle, T. M., & Turner, E. C. (2020). How butterflies keep their cool: Physical and ecological traits influence thermoregulatory ability and population trends. Journal of Animal Ecology, 89(11), 2440–2450. 10.1111/1365-2656.13319

Bowers, M. D., Brown, I. L., & Wheye, D. (1985). Bird predation as a selective agent in a butterfly population. Evolution, 39(1), 93–103. 10.2307/2408519

Brakefield, P. M., & Reitsma, N. (1991). Phenotypic plasticity, seasonal climate and the population biology of *Bicyclus* butterflies (Satyridae) in Malawi. Ecological Entomology, 16(3), 291–303. 10.1111/j.1365-2311.1991.tb00220.x

Breuker, C. J., Brakefield, P. M., & Gibbs, M. (2007). The association between wing morphology and dispersal is sex-specific in the glanville fritillary butterfly *Melitaea cinxia* (Lepidoptera: Nymphalidae). European Journal of Entomology, 104(3), 445–452. 10.14411/eje.2007.064

Büyükyilmaz, E., & Tseng, M. (2022) Developmental temperature predicts body size, flight, and pollen load in a widespread butterfly. Ecological Entomology 47(5):872–82. doi: 10.1111/een.13177

Cant, E. T., Smith, A. D., Reynolds, D. R., & Osborne, J. L. (2005). Tracking butterfly flight paths across the landscape with harmonic radar. Proceedings of the Royal Society B: Biological Sciences, 272(1565), 785–790. 10.1098/rspb.2004.3002

Chazot, N., Panara, S., Zilbermann, N., Blandin, P., Le Poul, Y., Cornette, R., Elias, M., & Debat, V. (2016). *Morpho* morphometrics: Shared ancestry and selection drive the evolution of wing size and shape in *Morpho* butterflies. Evolution, 70(1), 181–194. 10.1111/evo.12842

Cormont, A., Malinowska, A. H., Kostenko, O., Radchuk, V., Hemerik, L., WallisDeVries, M. F., & Verboom, J. (2011). Effect of local weather on butterfly flight behaviour, movement, and colonization: Significance for dispersal under climate change. Biodiversity and Conservation, 20(3), 483–503. 10.1007/s10531-010-9960-4

Danks, H. V. (2007). The elements of seasonal adaptations in insects. The Canadian Entomologist, 139(1), 1–44. 10.4039/n06-048

Dell’Aglio, D. D., Mena, S., Mauxion, R., McMillan, W. O., & Montgomery, S. H. (2022). Divergence in *Heliconius* flight behaviour is associated with local adaptation to different forest structures. Journal of Animal Ecology, 91(4), 727–737. 10.1111/1365-2656.13675

DeVries, P. J., Penz, C. M., & Hill, R. I. (2010). Vertical distribution, flight behaviour and evolution of wing morphology in *Morpho* butterflies. Journal of Animal Ecology, 79(5), 1077– 1085. 10.1111/j.1365-2656.2010.01710.x

Ellington, C. P. (1984). The aerodynamics of hovering insect flight. I. The quasi-steady analysis. Philosophical Transactions of the Royal Society of London. Series B, Biological Sciences, 305(1122), 1–15.

Evans, L. C., Oliver, T. H., Sims, I., Greenwell, M. P., Melero, Y., Watson, A., Townsend, F., & Walters, R. J. (2020). Behavioural modes in butterflies: Their implications for movement and searching behaviour. Animal Behaviour, 169, 23–33. 10.1016/j.anbehav.2020.09.001

Friberg, M., & Wiklund, C. (2019). Host preference variation cannot explain microhabitat differentiation among sympatric *Pieris napi* and *Pieris rapae* butterflies. Ecological Entomology, 44(4), 571–576. 10.1111/een.12728

Fric, Z., Klimova, M., & Konvicka, M. (2006). Mechanical design indicates differences in mobility among butterfly generations. Evolutionary Ecology Research, 8: 1511–1522.

Harrison, R. G. (1980). Dispersal polymorphisms in insects. Annual Review of Ecology and Systematics, 11, 95–118.

Hwang, Shaw-Yhi, Cheng-Hsiang Liu, and Tse-Chi Shen. 2008. Effects of plant nutrient availability and host plant species on the performance of two *Pieris* butterflies (Lepidoptera: Pieridae). Biochemical Systematics and Ecology 36(7):505–13. doi: 10.1016/j.bse.2008.03.001.

Jackson, B. E., Evangelista, D. J., Ray, D. D., & Hedrick, T. L. (2016). 3D for the people: Multi-camera motion capture in the field with consumer-grade cameras and open source software. Biology Open, 5(9), 1334–1342. 10.1242/bio.018713

Jantzen, B., & Eisner, T. (2008). Hindwings are unnecessary for flight but essential for execution of normal evasive flight in Lepidoptera. Proceedings of the National Academy of Sciences, 105(43), 16636–16640. 10.1073/pnas.0807223105

Josse, J., & Husson, F. (2016). missMDA: A package for handling missing values in multivariate data analysis. Journal of Statistical Software, 70(1), 1–31. 10.18637/jss.v070.i01

Kerr, N. Z., Wepprich, T., Grevstad, F. S., Dopman, E. B., Chew, F. S., & Crone, E. E. (2020). Developmental trap or demographic bonanza? Opposing consequences of earlier phenology in a changing climate for a multivoltine butterfly. Global Change Biology, 26(4), 2014–2027. 10.1111/gcb.14959

Kingsolver, J. G. (1985). Thermoregulatory significance of wing melanization in *Pieris* butterflies (Lepidoptera: Pieridae): physics, posture, and pattern. Oecologia, 66(4), 546–553. 10.1007/BF00379348

Kleckova, I., Konvicka, M., & Klecka, J. (2014). Thermoregulation and microhabitat use in mountain butterflies of the genus *Erebia*: Importance of fine-scale habitat heterogeneity. Journal of Thermal Biology, 41, 50–58. 10.1016/j.jtherbio.2014.02.002

Le Roy, C., Debat, V., & Llaurens, V. (2019). Adaptive evolution of butterfly wing shape: From morphology to behaviour. Biological Reviews, 94(4), 1261–1281. 10.1111/brv.12500

Le Roy, C., Roux, C., Authier, E., Parrinello, H., Bastide, H., Debat, V., & Llaurens, V. (2021). Convergent morphology and divergent phenology promote the coexistence of *Morpho* butterfly species. Nature Communications, 12(1), Article 1. 10.1038/s41467-021-27549-1

Lenth, R. (2022). emmeans: Estimated marginal means, aka least-squares mean (R package version 1.8.1-1) , https://CRAN.R-project.org/package=emmeans.

Losos, J, B., Creer, D. A., & Schulte II, J.A (2002). Cautionary comments on the measurement of maximum locomotor capabilities. Journal of Zoology 258 (1): 57–61.

Lyytinen, A., Brakefield, P. M., Lindström, L., & Mappes, J. (2004). Does predation maintain eyespot plasticity in *Bicyclus anynana*? Proceedings of the Royal Society of London. Series B: Biological Sciences, 271(1536), 279–283. 10.1098/rspb.2003.2571

Lyytinen, A., Alatalo, R. V., Lindström, L., & Mappes, J. (1999). Are European white butterflies aposematic? Evolutionary Ecology, 13(7), 709. 10.1023/A:1011081800202

Merckx, T., & Van Dyck, H. (2006). Landscape structure and phenotypic plasticity in flight morphology in the butterfly *Pararge aegeria*. Oikos, 113(2), 226–232. 10.1111/j.2006.0030-1299.14501.x

Moradinour, Z., Wiklund C., Miettinen M., Gérard M., & Baird E. (2023). Exposure to elevated temperature during development affects eclosion and morphology in the temperate *Pieris napi* butterfly (Lepidoptera: Pieridae). Journal of Thermal Biology 118:103721. doi: 10.1016/j.jtherbio.2023.103721.

Muijres, F. T., Elzinga, M. J., Melis, J. M., & Dickinson, M. H. (2014). Flies evade looming targets by executing rapid visually directed banked turns. Science, 344(6180), 172–177. 10.1126/science.1248955

Nijhout, H. F. (2003). Development and evolution of adaptive polyphenisms. Evolution & Development, 5(1), 9–18. 10.1046/j.1525-142X.2003.03003.x

Nylin, S. (1994). Seasonal plasticity and life-cycle adaptations in butterflies. In H. V. Danks (Ed.), Insect life-cycle polymorphism: Theory, evolution and ecological consequences for seasonality and diapause control (pp. 41–67). Springer Netherlands. 10.1007/978-94-017-1888-2_3

Ohsaki, N., & Sato, Y. (1999). The role of parasitoids in evolution of habitat and larval food plant preference by three *Pieris* butterflies. Population Ecology, 41(1), 107–119. 10.1007/PL00011975

Okamura, Y., Sato, A., Tsuzuki, N., Murakami, M., Heidel-Fischer, H., & Vogel, H. (2019). Molecular signatures of selection associated with host plant differences in *Pieris* butterflies. Molecular Ecology, 28(22), 4958–4970. 10.1111/mec.15268

R Core Team. (2022). R: A language and environment for statistical computing. R Foundation for Statistical Computing, Vienna, Austria. (4.2.1) [Computer software]. https://www.R-project.org/

Remmel, T., Tammaru, T., & Mägi, M. (2009). Seasonal mortality trends in tree-feeding insects: A field experiment. Ecological Entomology, 34(1), 98–106. 10.1111/j.1365-2311.2008.01044.x

Root, R. B., & Kareiva, P. M. (1984). The Search for resources by cabbage butterflies (*Pieris rapae*): Ecological consequences and adaptive significance of markovian movements in a patchy environment. Ecology, 65(1), 147–165. 10.2307/1939467

Ryan, S. F., Lombaert, E., Espeset, A., Vila, R., Talavera, G., Dincă, V., Doellman, M. M., Renshaw, M. A., Eng, M. W., Hornett, E. A., Li, Y., Pfrender, M. E., & Shoemaker, D. (2019). Global invasion history of the agricultural pest butterfly *Pieris rapae* revealed with genomics and citizen science. Proceedings of the National Academy of Sciences, 116(40), 20015–20024. 10.1073/pnas.1907492116

Sang, A., & Teder, T. (2011). Dragonflies cause spatial and temporal heterogeneity in habitat quality for butterflies. Insect Conservation and Diversity, 4(4), 257–264. 10.1111/j.1752-4598.2011.00134.x

Schneider, C. A., Rasband, W. S., & Eliceiri, K. W. (2012). NIH Image to ImageJ: 25 years of image analysis. Nature Methods, 9(7), Article 7. 10.1038/nmeth.2089

Shkurikhin, A. O., & Oslina, T. S. (2016). Seasonal variation of the forewing in polyvoltine whites *Pieris rapae* L. and *P. napi* L. (Lepidoptera: Pieridae) in the forest-steppe zone of the Southern Urals. Russian Journal of Ecology, 47(3), 296–301. 10.1134/S1067413616030115

Skórka, P., Nowicki, P., Kudłek, J., Pępkowska, A., Śliwińska, E., Witek, M., Settele, J., & Woyciechowski, M. (2013). Movements and flight morphology in the endangered Large Blue butterflies. Open Life Sciences, 8(7), 662–669. 10.2478/s11535-013-0190-5

Spieth, H., & Cordes, R. (2012). Geographic comparison of seasonal migration events of the large white butterfly, *Pieris brassicae*. Ecological Entomology, 37. 10.1111/j.1365-2311.2012.01385.x

Srygley, R. B. (1997). Locomotor mimicry in butterflies? The associations of positions of centres of mass among groups of mimetic, unprofitable prey. Philosophical Transactions of the Royal Society of London. Series B: Biological Sciences, 343(1304), 145–155. 10.1098/rstb.1994.0017

Srygley, R. B. (1999). Locomotor mimicry in *Heliconius* butterflies: Contrast analyses of flight morphology and kinematics. Philosophical Transactions of the Royal Society of London. Series B: Biological Sciences, 354(1380), 203–214. 10.1098/rstb.1999.0372

Tenger-Trolander, A., Julick, C. R., Lu, W., Green, D. A., Montooth, K. L., & Kronforst, M. R. (2023). Seasonal plasticity in morphology and metabolism differs between migratory North American and resident Costa Rican monarch butterflies. Ecology and Evolution, 13(2), e9796. 10.1002/ece3.9796

Thomas, C. D., Hill, J. K., & Lewis, O. T. (1998). Evolutionary consequences of habitat fragmentation in a localized butterfly. Journal of Animal Ecology, 67(3), 485–497. 10.1046/j.1365-2656.1998.00213.x

Tsuji, J. S., Kingsolver, J. G., & Watt, W. B. (1986). Thermal physiological ecology of *Colias* butterflies in flight. Oecologia, 69(2), 161–170.

Van Dyck, H., & Wiklund, C. (2002). Seasonal butterfly design: Morphological plasticity among three developmental pathways relative to sex, flight and thermoregulation. Journal of Evolutionary Biology, 15(2), 216–225. 10.1046/j.1420-9101.2002.00384.x

Van Rossum, G., & Drake Jr., F. L. (2009). Python 3 Reference Manual [Computer software]. Scotts Valley, CA: CreateSpace.

von Schmalensee, L., Caillault, P., Gunnarsdóttir, K. H., Gotthard, K., & Lehmann, P. (2023). Seasonal specialization drives divergent population dynamics in two closely related butterflies. Nature Communications, 14(1), Article 1. 10.1038/s41467-023-39359-8

Wickham, H., Hester, J., Chang, W., & Bryan, J. (2022). Devtools: Tools to make developing R packages easier [Computer software]. https://devtools.r-lib.org/

Wiklund, C., Nylin, S., & Forsberg, J. (1991). Sex-related variation in growth rate as a result of selection for large size and protandry in a bivoltine butterfly, Pieris napi. Oikos, 241-250.

Wootton, R.J. (1992). Functional morphology of insect wings. Annual Review of Entomology 37, 113–140.

Xue, B., & Leibler, S. (2018). Benefits of phenotypic plasticity for population growth in varying environments. Proceedings of the National Academy of Sciences, 115(50), 12745–12750. 10.1073/pnas.1813447115

Zvereva, E. L., & Kozlov, M. V. (2021). Seasonal variations in bird selection pressure on prey colouration. Oecologia, 196(4), 1017–1026. 10.1007/s00442-021-04994-9

